# Genome-scale metabolic modelling of *P. thermoglucosidasius* NCIMB 11955 reveals metabolic bottlenecks in anaerobic metabolism

**DOI:** 10.1101/2021.02.01.429138

**Authors:** Viviënne Mol, Martyn Bennett, Benjamín J. Sánchez, Beata K. Lisowska, Markus J. Herrgård, Alex Toftgaard Nielsen, David J. Leak, Nikolaus Sonnenschein

## Abstract

*Parageobacillus thermoglucosidasius* represents a thermophilic, facultative anaerobic bacterial chassis, with several desirable traits for metabolic engineering and industrial production. To further optimize strain productivity, a systems level understanding of its metabolism is needed, which can be facilitated by a genome-scale metabolic model. Here, we present *p-thermo*, the most complete, curated and validated genome-scale model (to date) of *Parageobacillus thermoglucosidasius* NCIMB 11955. It spans a total of 890 metabolites, 1175 reactions and 917 metabolic genes, forming an extensive knowledge base for *P. thermoglucosidasius* NCIMB 11955 metabolism. The model accurately predicts aerobic utilization of 22 carbon sources, and the predictive quality of internal fluxes was validated with previously published ^13^C-fluxomics data. In an application case, *p-thermo* was used to facilitate more in-depth analysis of reported metabolic engineering efforts, giving additional insight into fermentative metabolism. Finally, *p-thermo* was used to resolve a previously uncharacterised bottleneck in anaerobic metabolism, by identifying the minimal required supplemented nutrients (thiamin, biotin and iron(III)) needed to sustain anaerobic growth. This highlights the usefulness of *p-thermo* for guiding the generation of experimental hypotheses and for facilitating data-driven metabolic engineering, expanding the use of *P. thermoglucosidasius* as a high yield production platform.

## 1. Introduction

As the global transition away from petroleum-derived feedstocks continues, the need to produce commodity and fine chemicals using sustainable feedstocks has accelerated the interest in establishing microbial bioprocesses with lower environmental footprints^1–3^. The microbial ‘chassis’ organisms of these bioprocesses have been developed through modern metabolic engineering strategies. Such strategies have enabled the redirection of carbon flux in metabolic pathways of the corresponding microbes towards target products, in what are commonly termed ‘microbial cell factories’^4^.

Without an accurate picture of how cellular metabolism operates as a whole, metabolic engineering strategies can produce flux imbalances, resulting in the accumulation of carbon intermediates, metabolic bottlenecks and/or imbalances in the overall cellular redox ratio^5,6^. As a result, there can be large upfront costs in microbial strain engineering to ensure economically viable biochemical product yields^3,7^. To bolster traditional metabolic engineering efforts and help elucidate genotype-phenotype relationships, systems metabolic engineering aims to describe a more holistic representation of cellular metabolism though the integration of stoichiometric modelling and -omics data analyses^8^.

In particular, the advent of cheaper DNA sequencing has given rise to genome-scale metabolic models (GEMs), *in silico* reconstructions of the metabolic reaction networks of a given organism, derived from its annotated genome sequence^9^. In addition to operating as a knowledge base of metabolic information for a particular organism, GEMs can be used via constraint-based flux balance analysis to simulate carbon flux through metabolic reaction networks, enabling the rapid screening of metabolic behaviours under a range of environmental variables and biological contexts^10^. Through comprehensive *in silico* predictions of metabolic phentotypes under target conditions, GEMs can also identify potential cellular redox imbalances^11^ and metabolic bottlenecks and generate hypotheses for rational, targeted genetic modifications for improved performance^8^. GEMs can even guide the construction and optimization of carbon flux for either endogenous or novel heterologous microbial strain pathways towards high yields of desired products^12,13^.

*Parageobacillus thermoglucosidasius* NCIMB 11955 represents a Gram-positive, facultative anaerobic, thermophilic bacterial chassis with several advantageous traits for industrial bioprocesses when compared to many model bacterial chassis such as *Escherichia coli* and *Bacillus subtilis*^14,15^. Firstly, the thermophilicity of *Parageobacillus* spp. enables fermentations between 48-70°C^16,17^ at growth rates surpassing other thermophilic organisms^18^ and comparable to that of *E. coli*^19^. Compared to equivalent mesophilic fermentations, these process temperatures enable a reduction in both the cooling costs of large-scale exothermic fermentations, and a reduction in the risk of contamination from mesophilic microbes^8,20^. Furthermore, for industrial bioprocesses aiming for simultaneous saccharification and fermentation (SSF), the thermophilicity of *P. thermoglucosidasius* is complemented by a catabolic versatility. Through extracellular secretions of thermostable amylases^14^, xylanases^21–23^ and other hemicellulases^24–26^, *Parageobacillus* spp. are able to metabolise a wide range of C5 and C6 sugar monomers. Notably, they are able to transport then metabolize complex hemicellulosic^26^ and cellulosic^14^ polysaccharides derived from hydrolysates of lignocellulosic biomass, potentially reducing the reliance on externally supplied hydrolases involved in lignocellulosic pre-treatment.

A number of synthetic biology tools applicable *to P. thermoglucosidasius* have been devised including: shuttle vectors for reliable transformation^21,27^, chromosomal integration strategies^28^ promoter and RBS libraries to enable tuneable gene expression and validated reporter genes^29–32^. Such tools have enabled *P. thermoglucosidasius*, and genetically similar *(Para)geobacillus* spp., to be used in the production of fuels such as bioethanol^33,34^, isobutanol^35^ and hydrogen gas^36,37^ and also in fine chemicals including 2-3 butanediol^38,39^, riboflavin^40^ and isoprenoids^41^. *Parageobacillus* spp. and *Geobacillus* spp. have also been the source of thermostable variants of industrially useful proteases^42^, carboxyl esterases^43–45^, lipases^44,46^ along with a thermostable DNA polymerase I from *G. stearothermophilus* GIM1.543^47^.

In spite of these advances, (with the exception of natural end-products of glycolytic metabolism, such as ethanol) none of these engineered pathways have approached their potential maximum yields. In general, they have relied on natural flux to their metabolic precursors and its inherent control. The availability of a reliable GEM would enable a systems metabolic engineering approach of *P. thermoglucosidasius* to address the optimisation of flux through central metabolic pathways to balance the requirements of both production and growth. At present, only one publicly available GEM of a *P. thermoglucosidasius* exists, the related strain *P. thermoglucosidasius* C56-YS93 (denoted iGT736)^48^. While comprising 1159 reactions and 1163 metabolites, analysis of iGT736 using the GEM assessment tool Memote developed by Lieven et al.^49^ suggests that it currently lacks some fundamental features, including a biomass equation, transport reactions and stoichiometric balance (Supplementary File 1), preventing meaningful application for quantitative analysis. Additionally, a few examples exist of smaller central carbon metabolism scale models derived from experimental ^13^C isotopic tracer experiments. This includes models representing *P. thermoglucosidasius* M10EXG^7^ under aerobic and anaerobic growth conditions, and similar *Geobacillus* spp*. G. icigianus*^38^ and *Geobacillus* LC300^50^. However, they are less useful for illustrating the scale and complexity of whole cell metabolism.

The newly constructed genome-scale metabolic model *of P. thermoglucosidasius* NCIMB 11955 presented herein (named hereafter as *p-thermo*) represents 917 genes and comprises of 890 metabolites and 1175 reactions across two compartments: cytosolic and extracellular space (representing the medium). After iterative cycles of manual curation, model refinement and analysis with Memote^49^, *p-thermo* exhibits a 100% stoichiometric consistency, 100% charge balance and a 99.9% mass balance. It accurately captures experimentally determined utilization of 22 carbon sources using the sole input of measured production and consumption rates^51^ and is represented in the Systems Biology Markup Language (SBML)^52^ compliant format, making it compatible with commonly used constraint-based modeling software such as COBRApy^53^ and the COBRA Toolbox v3.0^54^ as well as more specialised software facilitating systems metabolic engineering^55,56^. Validation of the predictive quality of *p-thermo* under aerobic, oxygen limited and anaerobic conditions is demonstrated through mapping the resulting *in silico* fluxes to experimentally determined ^13^C-flux data obtained from ^13^C-isotopic labelling experiments of the genetically and metabolically similar *P. thermoglucosidasius* M10EXG strain^7^. The predictive power of *p-thermo* is further demonstrated through recapitulation of a metabolically engineered homoethanologenic strain of *P. thermoglucosidasius*^33^. Lastly, *p-thermo* was used to investigate the fundamental requirements and metabolic bottlenecks of *P. thermoglucosidasius* during anaerobic growth. The results establish a set of nutrients, namely biotin, thiamine and iron (III), that are required to support anaerobic growth of *P. thermoglucosidasius* NCIMB 11955 on a truly defined minimal media.

Currently, *p-thermo* represents the most complete, curated and experimentally validated genome-scale metabolic model for a *Parageobacillus* sp, and will be a foundational platform for guiding rational metabolic engineering strategies, -omic data integration, and strain optimization to further the potential of *P. thermoglucosidasius* NCIMB 11955 to operate as microbial chassis for sustainable bioprocesses.

## 2. Results

### 2.1 Model reconstruction

The presented genome-scale metabolic reconstruction of *P. thermoglucosidasius* NCIMB 11955 is based on genome sequencing by ERGO™ Integrated Genomics^57^ and Sheng *et al.*^58^. Genome annotation was performed through the ERGO™ Integrated Genomics suite^57^ and the RAST annotation server^59^, followed by gap filling with Pathway Booster^60^. The reconstruction was extensively manually curated using available literature and databases (KEGG, BRENDA, MetaCyc, MetaNetX and EC2PDB), according to benchmark approaches^61^. Detailed manual curation and refinement can be followed in Lisowska^14^ and in the GitHub repository.

This metabolic model consists of 890 metabolites, involved in a total of 1175 reactions, encoded for by 917 genes, across two compartments: cytosolic and extracellular space (representing the medium). Manual curation was critical to ensure complete consistency of the model (Supplementary report 1). Central carbon metabolism of the model resembles that of previously reported *(Para)geobacillus* spp^7,62^ (Figure 1A). Of all reactions, 9.3% are involved in transport or exchange, highlighting the flexibility of the strain to grow on various carbon sources (Figure 1B).

**Figure 1:**
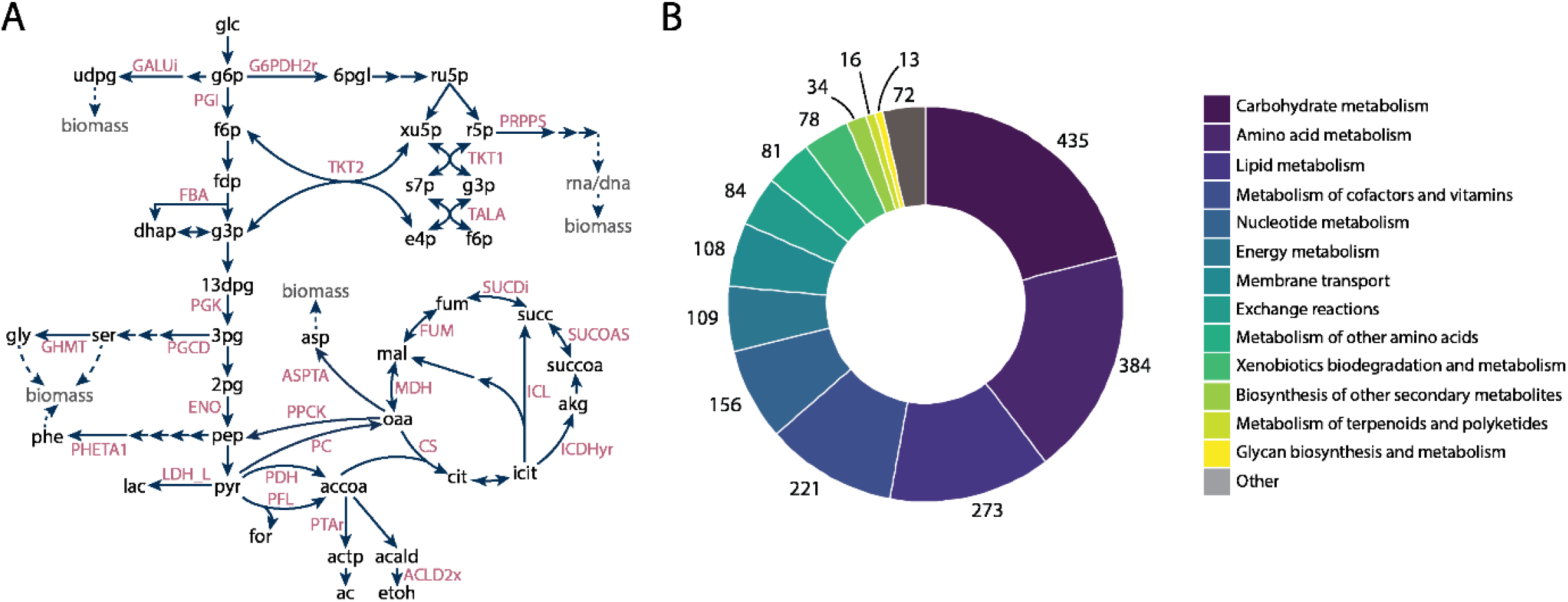
A) Central carbon metabolism map, with several reaction IDs highlighted. For a more detailed overview, see Supplementary File 3. B) The number of reactions in the model for several reaction class types. See section 5 for abbreviations used.

The model as well as scripts used in the reconstruction and manual curation are made publicly available through Github, at https://github.com/biosustain/p-thermo/releases/v1.0. The model is stored using the community-standard SMBL format (Level 3, FBC Version 2)^63^ and can additionally be accessed as Supplementary File 2.

### 2.2 Biomass composition and growth energetics

To capture biological growth in stoichiometric models, a demand reaction referred to as a biomass pseudo-reaction, was added. An overview of how the biomass pseudo-reaction was defined is explained in Materials & Methods, with the final reaction components and associated stoichiometry given in Supplementary Table 1. Energetic parameters were fitted from aerobically grown chemostat experiments. The energy required to maintain cellular homeostasis is reflected in the non-growth associated maintenance (NGAM) and was found to be 3.141 mmolATP/g_DW_h^−1^ in *p-thermo.* The growth associated maintenance, (GAM), was estimated as 152.3 mmol_ATP_/g_DW_ and reflects the energy needed for cell replication, including macromolecule synthesis. The contribution of polymerization energy, required for macromolecule synthesis, to the obtained GAM was estimated to be approximately 20% (Supplementary table 2); relatively low compared to previously reported mesophiles (30-40%)^64–67^. It was previously observed that thermophilic organisms tend to require higher levels of energy for growth and homeostasis at elevated temperatures and thus have a reduced growth efficiency, shown in the high maintenance estimated^18,68^. This trait of thermophiles makes them valuable hosts for bioproduction as it leads to higher production rates of catabolic products compared to other organisms.

### 2.3 Overview of metabolism

To provide a comprehensive overview, two pathway maps of the model were drawn using Escher^69^ corresponding to central carbon and amino acid metabolism (Supplementary Files 3 and 4) and deposited in the GitHub repository at p-thermo/maps. Traits specific to *Geobacillus* spp. and *P. thermoglucosidasius* NCIMB 11955, presented in the literature, were used to validate the model’s metabolism. Detailed step-by-step decisions that were made, can be followed in the GitHub repository at “p-thermo/notebooks”. As an example, in central carbon metabolism research has shown that *Geobacillius* spp., unlike many mesophilic *Bacillus* species, lack genes for a 6-phosphogluconolactonase (6PGL), responsible for part of the oxidative pentose phosphate pathway (PPP)^14^. Instead, the reaction can occur spontaneously and, at thermophilic temperatures, may be sufficiently rapid to maintain the requisite PPP flux^70^. The absence of 6PGL was captured in the model, but to reflect the active PPP pathway, a pseudo-reaction was added to allow the complete oxidative PPP to function.

*(Para)geobacillus* species are known to be capable of growth on a wide range of carbohydrates, and have been shown to secrete various polysaccharide degrading enzymes such as xylanases and other hemicellulose degrading enzymes^14,24–26^. To assess the metabolic capacity of the model, growth on various carbon sources was simulated (Figure 2). The choice of carbon sources was made based on what has previously been shown to allow aerobic growth of *P. thermoglucosidasius* NCIMB 11955^51^. Additionally, anaerobic growth on these substrates was computationally predicted. In both cases, carbon supply was normalized to 30 Cmols/gDWh, to accommodate different polymeric substrate forms being present in the data set. Initially, the model showed no aerobic growth on arbutin, salicin and rhamnose, due to dead-end metabolites being formed as side products in the first steps of their break down. Available literature was used to fill the gaps in the catabolic pathways, which enabled aerobic growth on all three carbon sources. Anaerobically, *in silico* growth on arbutin and salicin was unfeasible, as current knowledge suggests that their catabolism is oxygen dependent. Both arbutin and salicin are non-conventional carbon sources and are glycosides, composed of either a hydroquinone or salicyl alcohol functional group attached to glucose, respectively. It is known that metabolism of these glycosides occurs through splitting of the glycosidic bond, with the two functional groups being catabolized individually. With currently available knowledge, the further breakdown of the salicyl alcohol and hydroquinone functional groups is dependent on oxygen, deeming the *in silico* prediction of anaerobic growth unfeasible. As there is little knowledge about microbial catabolism of these carbon substrates, this hypothesis would warrant experimental validation.

**Figure 2:**
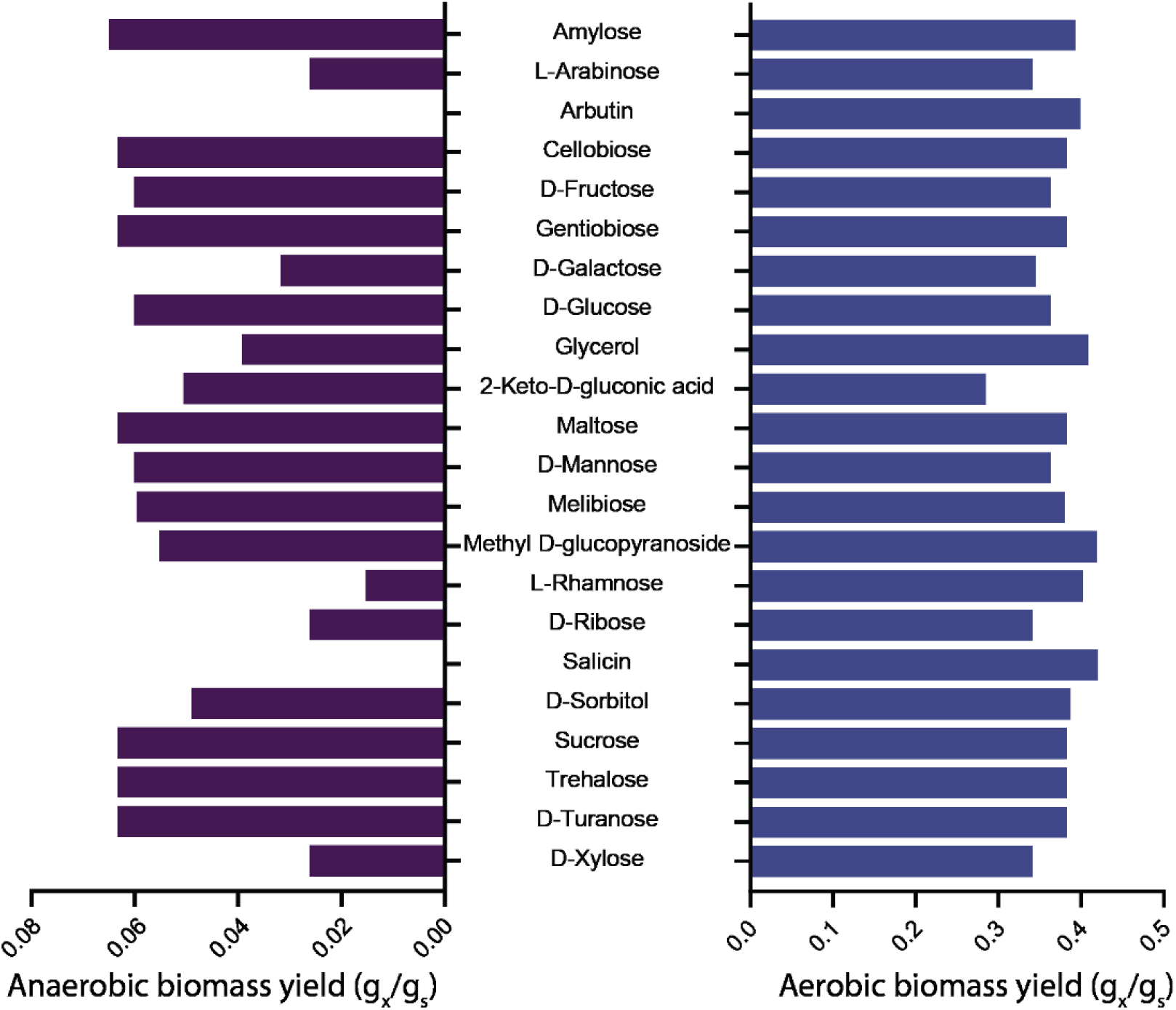
Anaerobic (left) and aerobic (right) predicted biomass yields for 22 different carbon sources, for which aerobic growth has been experimentally confirmed51. Carbon substrates were all supplied in the model at 30 Cmol/gDWh to account for differences in composition between the carbon sources.

### 2.4 Assessment of predictive power through ^13^C-flux fitting

Prior to using a genome-scale model for metabolic analyses or *ab initio* predictions, it is critical to validate its predictive power based on previously attained experimental data. This was done by analysis of how well simulated fluxes match known flux distributions. ^13^C-isotopic labelling is a standard tool used to elucidate intracellular fluxes in central carbon metabolism, through extensive experimental work and data analysis. Flux variability analysis (FVA) is an *in silico* approach that can allow *ab initio* analysis of metabolism without the need for laborious experimental data71. Comparing the two data types can give insights into metabolism and allow the generation of hypotheses for metabolic engineering purposes.

To do so, ^13^C-flux data from *P. thermoglucosidasius* M10EXG subject to varying oxygen conditions was used to qualitatively assess the predictive quality of *p-thermo*^7^. Whole proteome analysis (on a sequence basis) of the P*. thermoglucosidaius* M10EXG and NCIMB 11955 strains shows that the ORFs between the two strains are highly similar (Supplementary figure 1, Supplementary table 3). Specifically considering metabolic genes that would be captured as reactions in a metabolic model, there are only 11 and 12 unique reactions in *P. thermoglucosidasius* NCIMB 11955 and *P. thermoglucosidasius* M10EXG respectively (Supplementary tables 3, 4 and 5). Therefore, based on the overall metabolic similarity between the two strains, we assume that the ^13^C-flux data from *P. thermoglucosidasius* M10EXG can be utilized for a qualitative assessment.

In order to test if the model can predict intracellular fluxes close to the ^13^C-flux data, the measured production and consumption rates were fixed in the model as exchange rates, and internal fluxes were predicted in a sensitivity analysis with FVA. Parsimonious enzyme usage flux balance analysis (pFBA) which has previously been shown to predict fluxes that correlate with experimental measurements, was also performed^72^. The *in silico* fluxes and pFBA results were mapped to experimentally determined fluxes in aerobic, oxygen limited and anaerobic conditions (Figure 3A, B and C respectively). pFBA showed good correlation to the measured data for each condition (Supplementary Figure 2), and together with FVA showed accurate predictions of the internal central carbon fluxes (Figure 3A, B, and C). Predicted biomass yields (Figure 3D) and oxygen consumption rates (Supplementary Figure 3) were adequately predicted as well. These analyses validate the predictive quality of the created model and highlight the power of using metabolic models for understanding intracellular fluxes when only extracellular consumption or production rates are available.

**Figure 3:**
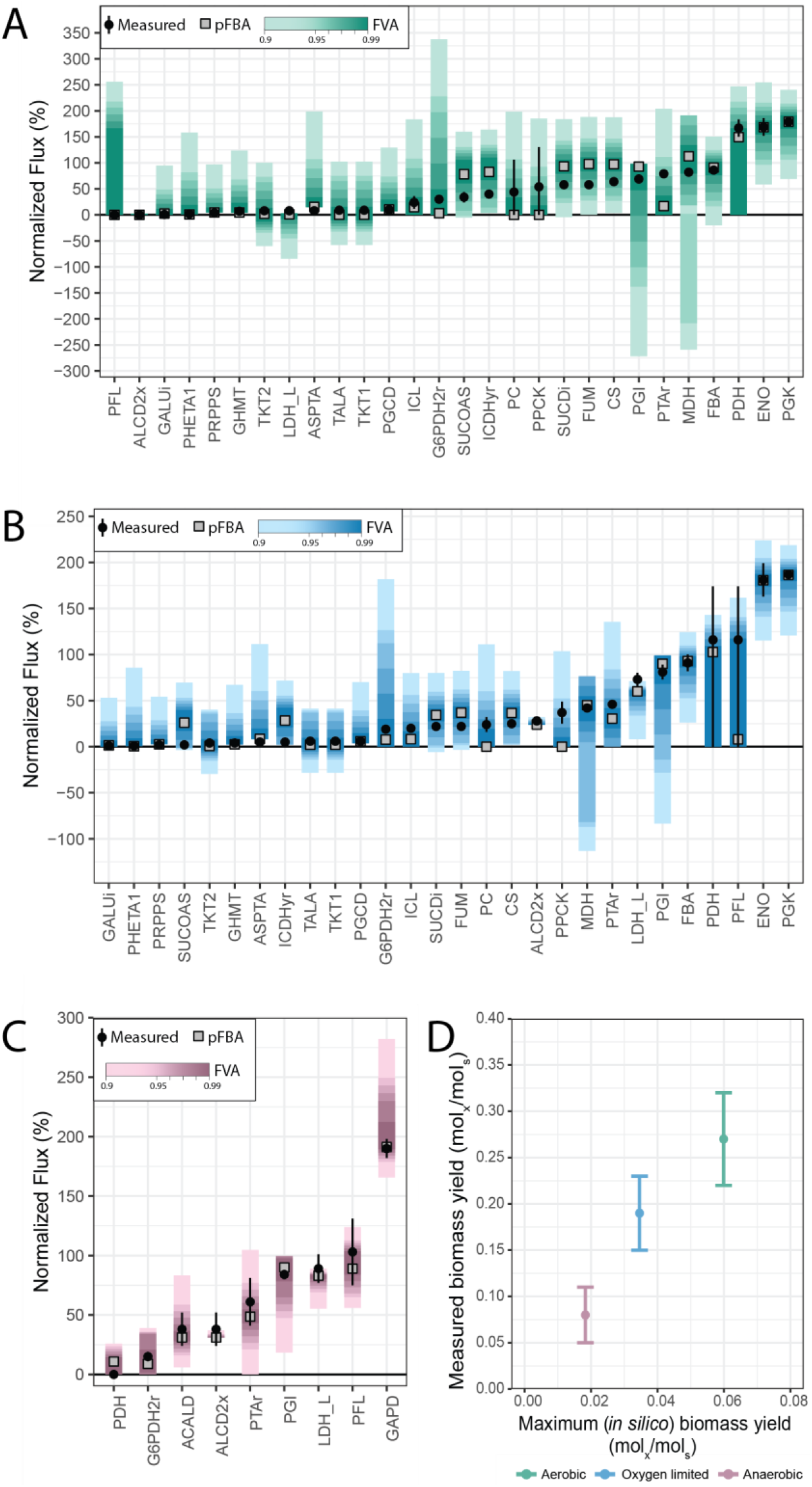
Results of fixing experimentally measured exchange rates and predicting intracellular flux distributions7 in aerobic (A), oxygen limited (B) and anaerobic (C) conditions, normalized to the glucose uptake rate. Figure 1A shows the stoichiometry of all the reactions shown on the x-axis. See Section 5 for abbreviations. FVA sensitivity analysis is shown in line ranges. Predicted and measured maximum biomass yields, for a FVA threshold set at 99% of optimum biomass, are shown in (D).

### 2.5 Recapitulating & interpreting knockout physiology

To evaluate the utility of *p-thermo* for metabolic engineering applications, we recreated previously reported homoethanologenic mutants of *P. thermoglucosidasius* NCIMB 11955 *in silico.* Cripps *et al*^33^ engineered lactate dehydrogenase (*ldh*) and pyruvate formate lyase (*pfl*) knockouts *in vivo*, and supplemented their ethanol yields with an upregulation of pyruvate dehydrogenase expression (PDH_up_). Using *p-thermo*, the wild type (WT), Δ*ldh* and Δ*ldh*Δ*pfl*(PDH_up_) strains were recreated, as stoichiometric modeling cannot distinguish between upregulated expression levels (i.e. between Δ*ldh*Δ*pfl* and Δ*ldh*Δ*pfl* PDH_up_). Exchange rates of the main fermentation metabolites were predicted using *p-thermo* and their accuracy evaluated based on measured data (Figure 4). In performing the analysis, two distinct thresholds for Flux Variability Analysis (FVA) were selected, 95% and 99% of optimum biomass production^71^, to assess the flexibility of exchange rates to the simulated conditions.

**Figure 4:**
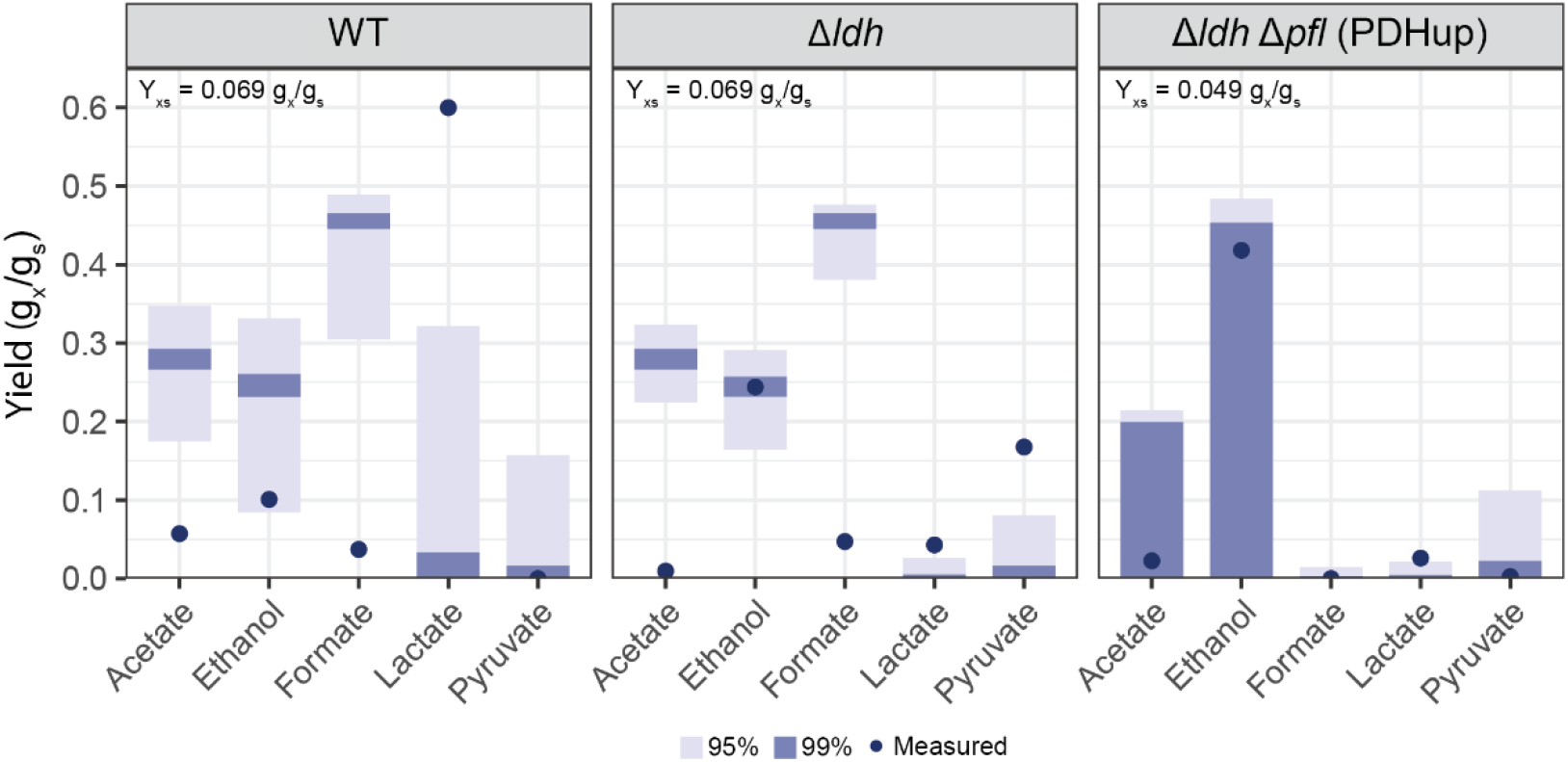
Comparison of *in silico* predictions of fermentation product yields in three engineered strains with experimentally determined data from Cripps *et al.*^33^, when solely the carbon uptake rate and knockouts were fixed in the model. Yield (g_x_/g_s_) is shown for predicted and measured exchange rates. Each panel highlights a different strain: wild type (WT), Δ*ldh* and Δ*ldh*Δ*pfl* (PDHup). Two varying thresholds for FVA were run: 95% and 99% of the optimum biomass production. The *in silico* predicted biomass yield (Y_xs_) for 99% of the optimum biomass production is shown for each condition.

The performed simulations show a substantial discrepancy between predicted and measured yields in the WT and Δ*ldh* strains, whereas simulations tightly match the measured yields in the Δ*ldh*Δ*pfl* PDH_up_ strain. Still, the mismatch between the experimental and *in silico* data, *p-thermo* can be used to understand metabolic branch points. The main discrepancy observed lies in the lactate and formate yields for the WT and Δ*ldh* strains, which can be traced to the cellular decision of what to do with the synthesized pyruvate (Figure 4). In this regard, there are three options: 1) conversion into lactate by lactate dehydrogenase (LDH), 2) anaerobic conversion into acetyl-CoA by pyruvate formate lyase (PFL) or 3) aerobic conversion into acetyl-CoA by pyruvate dehydrogenase (PDH). In both the WT and Δ*ldh* strain, *p-thermo* showed flux from pyruvate to acetyl-CoA to be exclusively carried through PFL, fitting with experimental expectations under anaerobic conditions due to high [NADH]^73^. Additionally, the conversion of pyruvate into acetate and ethanol results in one additional ATP per glucose, compared to converting pyruvate into lactate^74^. Therefore, from a stoichiometric perspective, *p-thermo* predicts this to be the most optimal pathway for growth, explaining the high concentrations of formate, ethanol and acetate predicted in the simulation.

However, this was not observed in the experimental yields, presumably because of subtle differences in dissolved oxygen availability in the experimental setup that influence multiple levels of regulation *in vivo*, intrinsically not accurately captured by stoichiometric models. In the experimental dataset, undefined oxygen limited conditions were used in which a gradual decline in available dissolved oxygen concentration would have occurred during growth, whereas simulations were performed anaerobically. Under oxygen-limited conditions, PDH is expressed in the wild type *P. thermoglucosidasius*^33^, where PFL is typically only active under completely anaerobic conditions^75^. The transition of physiological states in response to decreasing oxygen availability results in excess NADH and creates a redox imbalance in the cell which is alleviated through production of lactate as the production of formate by PFL is restricted. This could explain the discrepancy between the experimentally measured low formate and high lactate production in the WT strain and the prediction by *p-thermo*. In the LDH knockout at low dissolved oxygen conditions, which prevents PFL activity, PDH instead predominantly carries flux to acetyl-CoA. In this instance, in order to maintain cellular redox balance, the Δ*ldh* cells increase the produced ethanol/acetate ratio. This picture highlights the complexity of cellular and enzymatic regulation that is poorly captured in stoichiometric models, as well as the difficulty in simulating uncontrolled environments accurately.

However, the performed simulations can still be used to visualize and understand the burden that lactate production can have on cellular growth. The inability to induce PFL at moderate levels of oxygen limitation, puts a larger reliance on fermentative metabolism to lactate, providing less energy. We used *p-thermo* to investigate the possible impact this has. First, all measured exchange rates for the three strains were fitted to the model and used in subsequent determination of predicted biomass yields. This showed that the model is physiologically capable of capturing the measured data, albeit with a lower predicted biomass yield than was experimentally measured, suggesting that stoichiometrically sub-optimal fermentation pathways were active *in vivo* (Supplementary Figure 4A). Finally, the effect of increasing lactate production on biomass yield was computed, showing the energetic loss that occurs from lactate production (Supplementary Figure 4B). Overall, this highlights the importance of complex regulation dictating metabolism, over pure stoichiometric optima per se.

### 2.6 Genome-scale metabolic modeling allows the elucidation of metabolic bottlenecks

The availability of a comprehensive GEM can also facilitate the elucidation of metabolic bottlenecks and identification and optimization of chemically defined growth media^76^. Thus, *p-thermo* was used to help resolve known issues of the anaerobic metabolic physiology of *P. thermoglucosidasius.* Although it is clearly capable of classical mixed acid fermentation and shows elements of a regulated aerobic-anaerobic switch as revealed by transcriptomic analysis^77^ (although the aerobic respiratory electron transport chain remained active in an oxygen-scavenging state under fermentative conditions) it has long been known that growth under anaerobic conditions requires additional growth supplements to those under aerobic conditions. Typically, this was resolved by supplementation with a small amount of oxygen or yeast extract^14,33,51^. Therefore, here we used simulations of *p-thermo* to find a minimal set of defined nutrients that can achieve anaerobic growth of *P. thermoglucosidasius*.

As a first observation, when fed true minimal, anaerobic medium, the model predicted no growth, in accordance with experimental observations. However, fermentative energy generation was observed, which highlights that oxygen requirement comes from critical secondary metabolites or cofactors that cannot be synthesized anaerobically, which is corroborated by previous observations^14^. By minimizing the oxygen uptake in the model, a critical reaction set requiring oxygen was generated (Table 1). This analysis highlighted a complex combination of components that cannot be synthesized anaerobically: thiamine, biotin, folate, vitamin B12, spermine, spermidine and hemin. Additionally, iron(III) must be available in the medium to allow porphyrin biosynthesis.

**Table 1:**
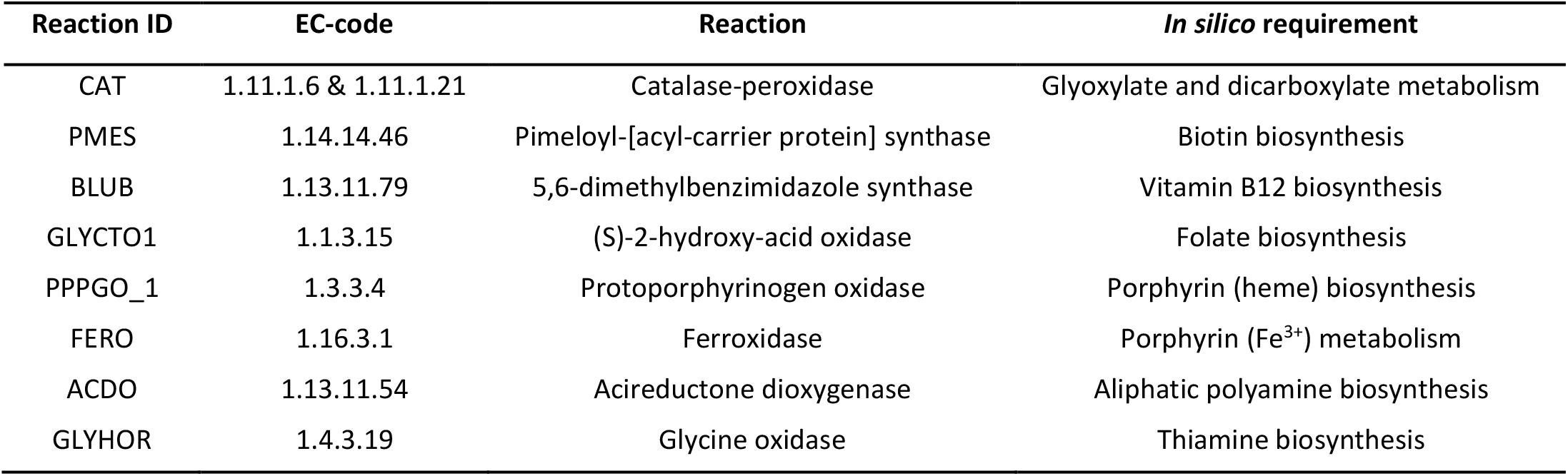
Overview of critical reactions that require oxygen to allow growth in p-thermo.

| Reaction ID | EC-code | Reaction | *In silico* requirement |
| --- | --- | --- | --- |
| CAT | 1.11.1.6 & 1.11.1.21 | Catalase-peroxidase | Glyoxylate and dicarboxylate metabolism |
| PMES | 1.14.14.46 | Pimeloyl-[acyl-carrier protein] synthase | Biotin biosynthesis |
| BLUB | 1.13.11.79 | 5,6-dimethylbenzimidazole synthase | Vitamin B12 biosynthesis |
| GLYCTO1 | 1.1.3.15 | (S)-2-hydroxy-acid oxidase | Folate biosynthesis |
| PPPGO_1 | 1.3.3.4 | Protoporphyrinogen oxidase | Porphyrin (heme) biosynthesis |
| FERO | 1.16.3.1 | Ferroxidase | Porphyrin (Fe<sup>3+</sup>) metabolism |
| ACDO | 1.13.11.54 | Acireductone dioxygenase | Aliphatic polyamine biosynthesis |
| GLYHOR | 1.4.3.19 | Glycine oxidase | Thiamine biosynthesis |

As expected, *in silico* supplementation of these components rescued anaerobic growth, providing a combination of candidates for experimental validation. The simulated essential components were experimentally added together in trace amounts to form a supplementation mix (see Materials and Methods); to assess if it would allow anaerobic growth. This was compared to Wolfe’s vitamin solution, a commonly used mix of vitamins in base thermophilic minimal medium (TMM)^31,78^. It should be noted that Wolfe’s vitamin solution contains thiamin, biotin, folate and vitamin B12, amongst other nutrients, and that TMM contains trace amounts of iron(III). To uncover the minimal sets of components needed to rescue anaerobic growth, eight different conditions were tested, all composed of base TMM with 10 g/l glucose: 1) no added nutrients, 2) 0.2% yeast extract, 3) biotin, 4) thiamin, 5) biotin and thiamin, 6) Wolfe’s vitamins, 7) Supplementation mix and 8) Wolfe’s vitamins plus the unique components of the supplementation mix (spermine, spermidine and heme) (Figure 5, Supplementary figure 5).

**Figure 5:**
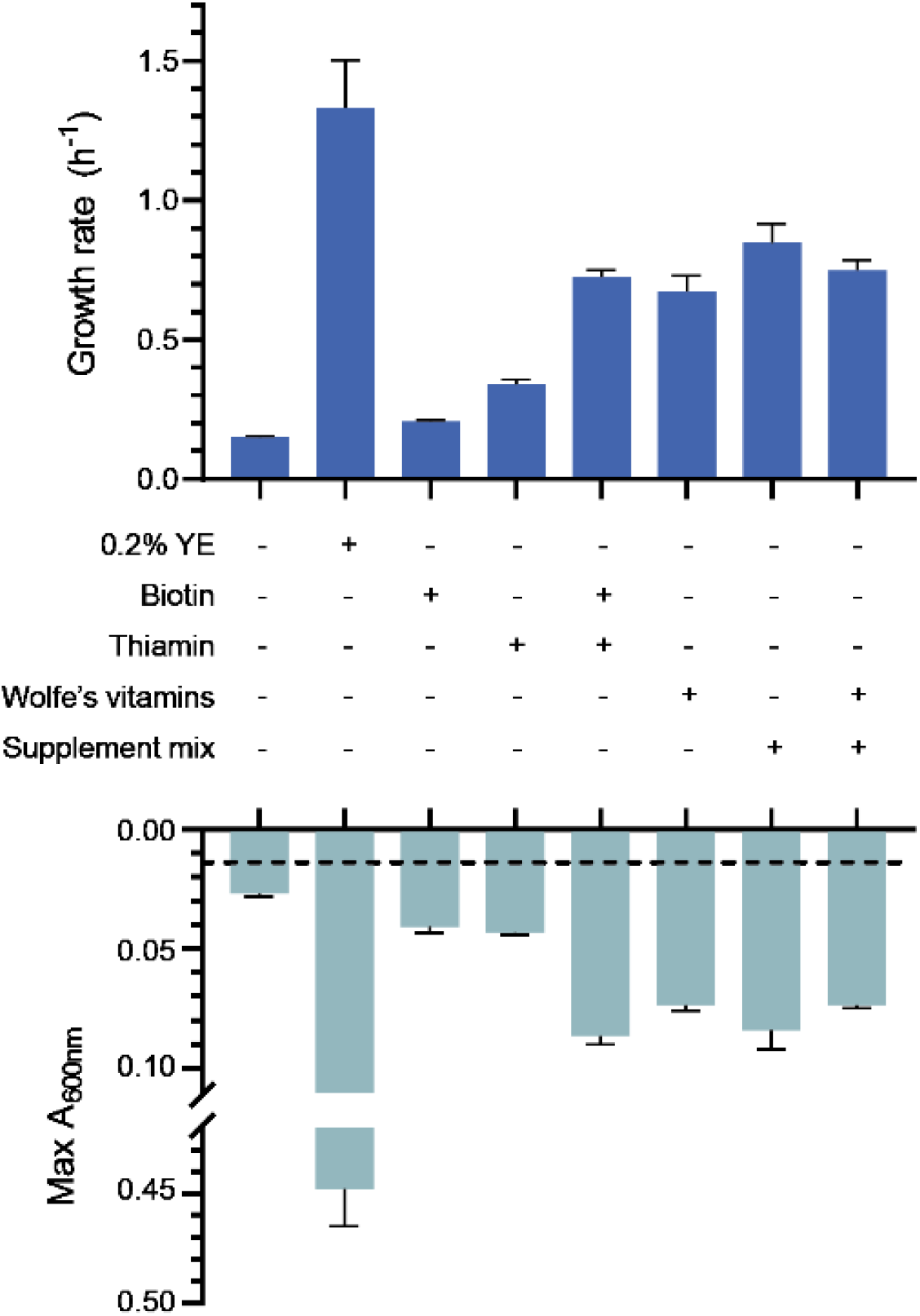
Experimental growth rates calculated and maximum observed absorbance values when *P. thermoglucosidasius* NCIMB 11955 was grown anaerobically in a microtiter plate reader in TMM base medium, supplemented with various nutrients, as indicated. Dashed line indicates inoculation absorbance, when an inoculation optical density of 0.05 was used.

Experimental observations suggested that a combination of thiamin, biotin and iron(III) were the minimal required supplementation set needed to sustain anaerobic growth, as no difference was observed when additional defined supplementation was added (Figure 5). Yeast extract also contains significant amounts of amino acids and other components and so provides an additional growth advantage, as expected. However, this highlights a discrepancy with the model predictions, as a larger minimal supplementation set was originally predicted (Table 1). Finally, as expected, base TMM can support aerobic growth, at a maximum rate of 0.267± 0.021 h^−1^ (Supplementary figure 5A), confirming the synthesis of the critical components in the presence of oxygen.

There are several reasons that can explain the differences between the *in silico* and experimentally determined minimal supplementation set. The incomplete understanding of thermophilic life introduces additional levels of complexity that are typically not captured by automatic annotation pipelines dependent on predominantly mesophilic datasets^59,79^ leading to errors in the annotation of thermophilic traits^80^. For example, genomes of thermophilic organisms show a correlation with higher G/C content, less intergenic regions and a higher functional stability (reflected by the lower ratio of non-synonymous to synonymous substitutions over time)^81–83^. Additionally, thermostable proteins can have significantly altered structure compared to their mesophilic counterparts performing the same reaction, confounding homology-based annotation^84^. Through the observed discrepancies, we can unveil additional insights of anaerobic metabolism of *P. thermoglucosidasius.*

In the first place, the *in silico* dependence on vitamin B12 highlights the inaccuracy of annotation pipelines. Vitamin B12 synthesis is classically divided into two routes: canonical (aerobic) and non-canonical (anaerobic)^85^. Although the genome annotation of *P. thermoglucosidasius* NCIMB 11955 reveals parts of either pathway, neither is complete. It has been proposed that possibly a novel, blended pathway may be present; however, this may arise from incorrect annotations based on lacking knowledge of thermophilic vitamin B12 biosynthesis genes^51,86,87^. The possibility to grow without vitamin B12 supplementation does highlight both an aerobic and anaerobic functional pathway in *P. thermoglucosidasius* NCIMB 11955. To further understand the *de novo* biosynthesis of vitamin B12, experimental validation would be required.

In *p-thermo*, the *in silico* oxygen requirement for spermine and spermidine biosynthesis comes from the downstream recycling of a biosynthetic by-product: 5’-methylthioadenosine (5-MTA). 5-MTA recycling is also important in a novel, oxygen independent MTA-isoprenoid shunt, involved in the methionine salvage pathway^88^. This pathway has been characterized in *Rhodospirillum rubrum* and orthology analysis highlights the possible presence of parts of this pathway in various facultative anaerobic *Bacillus* spp^89^. This presents the possibility of an alternate 5-MTA recycling pathway, explaining the independence of anaerobic growth to spermine or spermidine addition.

Similarly to spermine and spermidine, the oxygen requirement in the *in silico* folate biosynthesis pathway stems from the formation of glycolaldehyde as a side product, which is further oxidized to glyoxylate. The *in silico* oxidase responsible for glyoxylate formation requires oxygen (EC 1.1.3.15). However, reports show that *Moorella thermoacetica*, a thermophilic obligate anaerobe, can grow on glycolate through the formation of glyoxylate, highlighting the possibility for a (to date) unknown alternate electron acceptor^90,91^

Finally, heme is suggested to be synthesized in *P. thermoglucosidasius* from glycine using a 5-aminolevulinic acid synthase^51^ and notably using an oxygen-dependent protoporphyrinogen oxidase. Both *E. coli* and *B. subtilis* have an oxygen independent coproporphyrinogen-III oxidase (hemN), known to be responsible for anaerobic heme biosynthesis, using other electron acceptors such as fumarate, or nitrate over oxygen^92–95^. While the current genome annotation of *P. thermoglucosidasius* NCIMB 11955^58^ suggests that only the oxygen dependent path is present, the data presented herein suggest that supplementation with hemin is not required for growth (Figure 5). One possible explanation for this discrepancy can be found when performing a tBLASTn with the *B. subtilis* hemN (NCBI accession CAB61616) against the *P. thermoglucosidasius* NCIMB 11955 genome. This highlighted a significant hit (CP016622 region 3448674..3449762, 52% identity, E-value: 10^−115^). This suggests the possibility that some form of this oxygen independent heme biosynthesis route could also be present highlighting the need to better understand the metabolism of non-model organism chassis.

This identification of the minimal, defined anaerobic medium highlights how GEMs can be used to facilitate experimental hypotheses, where previous hypotheses have failed. The result, a defined minimal anaerobic medium is valuable for further investigation into anaerobic metabolism through ^13^C-characterization studies, where defined media are critical. Additionally, this identification of a series of components which support anaerobic growth of *P. thermoglucosidasius* at a minimal medium level can further help inform the development of industrial growth media for other microbial chassis used in anaerobic bioprocesses improving growth and chemical product yields.

## 3. Discussion

*Parageobacillus* spp. represent valuable microbial chassis for metabolic engineering and fermentative bioproduction. Many advantages derive from their thermophilic character, with additional advantages coming from species specific traits. However, to further develop *Parageobacillus* spp. into fully optimized microbial cell factories, additional in-depth and systems level understanding of metabolism is required, for which omic analyses and genome scale metabolic models are critical. Currently, various automatic pipelines exist for generating metabolic models on the sole basis of a genome sequence. Yet, for thermophilic organisms, significant faults resulting from automatic annotation pipelines are evident, as these are based on predominantly mesophilic datasets. Thermophilic genomes show different characteristics, the effect of which on metabolism is still poorly understood, making translation into a predicted function difficult. Thus, significant manual curation is needed in the generation of GEMs for thermophilic organisms, which is limited by the availability of knowledge on thermophilic metabolism. To increase the understanding of genotype-phenotype relationships in thermophilic hosts, the availability of a GEM acts as a considerable step facilitating systems level studies. As a result, with more knowledge arising, iterative rounds of model improvement are possible.

Therefore, in this work, we developed *p-thermo*, to date the most complete, curated and validated genome-scale metabolic model for a facultative anaerobic *Parageobacillus sp*. In it, genomic and biochemical knowledge were combined into a single powerful knowledge base, providing a critical tool for data-driven metabolic engineering, -omic data integration, process design and optimization. The model accurately captured substrate usage *in silico*, showing the metabolic flexibility of the strain for production with alternative carbon sources (Lisowska, 2016). Furthermore, ^13^C-isotopic data verified the quality of *p-thermo* for predicting central internal fluxes of the model, when solely production and consumption rates are measured, a common practice when evaluating metabolic engineering designs.

Going beyond validation, *p-thermo* was used to provide more in-depth analysis of previously reported metabolic engineering approaches^33^. Initial *p-thermo* simulations did not completely match experimental data, as stoichiometric models are incapable of capturing complex levels of regulation that play a dominant role in *in vivo* metabolism. Nonetheless, *p-thermo* was used to investigate the pyruvate branch point in central metabolism and allowed additional insights into metabolic flux distributions in various genetic backgrounds. With *p-thermo* further insights into metabolism can be gained, allowing improved targeted metabolic engineering in subsequent designs.

Finally, we used *p-thermo* to generate hypothesis driven experiments to alleviate a bottleneck in anaerobic metabolism, where previous experimental design was unsuccessful^14^. Doing so gave fundamental insights into the metabolism of *P. thermoglucosidasius*, and also showed that significant knowledge gaps still exist. This analysis, in combination with the ^13^C-based verification, highlights an additional obstacle in working with thermophilic GEMs, where annotation pipelines are less precise: information on central carbon metabolism can be inferred with relative accuracy, where peripheral metabolic pathways are not significantly understood and requires further systems-level investigation.

Overall, *p-thermo*, together with other systems level and omics based approaches, act as a tool to improve our understanding of genotype-phenotype relationships. Genome-scale metabolic models are in this way a critical part of an iterative cycle, and are essential to the use and efficacy of thermophilic hosts for metabolic engineering and industrial bioproduction.

## 4. Materials & Methods

### 4.1 Model construction & curation

Genome sequencing of *Parageobacillus thermoglucosidasius* NCIMB 11955 was initially performed by ERGOTM Integrated Genomics^57^ (funded by TMO Renewables Ltd) and subsequently updated using the published *P*. *thermoglucosidasius* NCIMB 11955 genome sequence^58^ (NCBI accession CP016622[chromosome], CP016623[pNCI001], and CP016624[pNCI002]). Genome annotation was performed through the ERGOTM Integrated Genomics suite^57^ and the RAST server^96^. Pathway Booster was used for gap filling, resolving gaps through comparisons with evolutionarily-related genomes^60^. Upon base construction of the model, further manual curation was done following standard procedures^61^. To do so, missing information was primarily obtained from literature and using various databases: BRENDA, EC2PDB, KEGG, MetaNetX or MetaCyc^51^. Whenever information on *P. thermoglucosidasius* was lacking, available references from other (*Para)geobacillus* spp or *Bacillus* spp were added. All further manual curation and refinement can be found in the GitHub repository. Model improvement was ensured by running Memote^49^ after each modification.

### 4.2 Biomass composition and growth energetics

To model growth, a biomass pseudo-reaction was added to the model. The reaction pools metabolites needed for growth into a biomass metabolite. Base biomass composition was previously determined experimentally according to reported practices^9,51^. Lipid composition was obtained from previous reports^7^, and was incorporated into the model according to a restrictive approach^97^, in which a determined acyl chain length is assumed for all lipid species. Further fine-tuning of biomass composition was performed based on available enzymatic and metabolic requirements of the strain, with case-by-case justification given in the GitHub repository. Critical metabolites known to be required for catabolic functions of essential enzymes, such as heme, were added at trace stoichiometries based on knowledge from related organisms and scaled to ensure that all biomass components added up to 1 g/g_DW_^98^.

Growth energetics (ATP cost of growth-associated maintenance and ATP requirement for non-growth associated maintenance) were estimated by minimizing the prediction error of the specific substrate consumption rate and the specific growth rate of glucose fed, aerobic chemostats^51,61^. The P/O ratios were obtained from the given data for *B. subtilis*^99^. The contribution of polymerization of each metabolite type to the total growth associated maintenance was estimated based on previously reported polymerization energies^64^.

### 4.3 Transport reactions

The model has two compartments: extracellular and intracellular. Transport reactions were inferred from genome annotations and homology to known transporters. Additionally, knowledge about growth on various substrates was used to validate the presence of the corresponding transporters.

### 4.4 Stoichiometric modeling & applied constraints

In traditional flux balance analysis^10^, reaction stoichiometries are converted into a stoichiometric matrix (**S**), with *m* × *n* dimensions, where *m* represents the various metabolites and *n* represents the number of reactions. Coefficients in the matrix are either positive or negative, reflecting production and consumption, respectively. Stoichiometric modeling works under the assumption of a pseudo-steady-state, represented as:

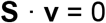

Where the vector **v** contains the fluxes of all reactions, given in units of mmol/g_DW_h. As there are more metabolites than reactions, to solve this underdetermined system, linear programming is used by formulating an objective function (*z*), per default set as biomass accumulation. Reversibility of reactions is set based on thermodynamic prediction and a default medium is defined; in the case of *p-thermo*, minimal medium, with D-glucose as default carbon source is used.

Quantification of metabolic fluxes was performed using flux variability analysis (FVA)^71^. When running FVA, a threshold below the optimum is used to represent the metabolic freedom that is given to a model. In this study, a sensitivity analysis was run with FVA thresholds from 90 to 99%, to evaluate at what sensitivity level the model better matches the experimental data. Additionally, parsimonious flux balance analysis (pFBA) was run, by conducting a bilevel linear programming optimization that computes the optimum (growth) solution of the network, whilst minimizing the sum of all fluxes^72^. In doing so, this optimization predicts the most stoichiometrically efficient pathway set, and captures the maximum biomass per unit flux objective that has previously been described to be well supported by proteomic and transcriptomic data^72,100^.

### 4.5 Genome comparison

For an unbiased genome comparison, the *P. thermoglucosidasius* NCIMB 11955 genome was obtained from NCBI (Accession: CP016622), and the *P. thermoglucosidasius* M10EXG genome was obtained from the Integrated Microbial Genome database (ID 2501416905). Genome annotation was performed using RASTk^96^, after which the two proteomes were compared through blast bi-directional best hits to create a homology matrix between the strains, based on a published pipeline^101^. To filter for metabolic genes, any ORF associated to a predicted EC code was considered metabolic. The exact workflow can be followed in the GitHub repository.

### 4.6 Experimental procedures

The *P. thermoglucosidasius* NCIMB 11955 (DSM2542) strain was obtained from DSMZ^102^. The strain was grown in either SPY medium or base thermophile minimal medium (TMM), modified from Fong *et al.*^78^. SPY was used for a first preculture, and contains per liter, 16g soy peptone, 10 g yeast extract and 5 g NaCl, adjusted to pH 6.8. Base TMM contains, per liter: 930 ml Six salts solution (SSS), 40 ml 1M MOPS (pH 8.2), 10 ml 1 mM FeSO_4_ in 0.4 M tricine, 10 ml 0.132 M K_2_HPO_4_, 10 ml 0.953 M NH_4_Cl, 0.5 ml 1M CaCl_2_ and trace element solution, adjusted to a final pH of 6.8. SSS contains, per 930 ml: 4.6 g NaCl, 1.35 g Na_2_SO_4_, 0.23 g KCl, 0.037 g KBr, 1.72 g MgCl_2_·6 H_2_O and 0.83g NaNO_3_. The trace element solution contained, per liter, 1g FeCl_3_·6 H_2_O, 0.18 g ZnSO_4_·7 H_2_O, 0.12 g CuCl_2_ ·2 H_2_O, 0.12 g MnSO_4_·H_2_O and 0.18 g CoCl_2_·6 H_2_O. D-glucose to a final concentration of 10g/L was added to the base TMM.

When indicated, the base TMM was supplemented with one of the following, to the indicated final concentrations: 0.2% (w/v) yeast extract, 2 μg/L biotin, 5 μg/L thiamine-HCl, 1x Wolfe’s vitamins, or 1x supplementation mix. 1000x Wolfe’s vitamins consist of, per liter, 10 mg pyridoxine hydrochloride, 5.0 mg thiamine-HCl, 5.0 mg riboflavin, 5.0 mg nicotinic acid, 5.0 mg calcium D-(+)-pantothenate, 5.0 mg p-aminobenzoic acid, 5.0 mg thioctic acid, 2.0 mg biotin, 2.0 mg folic acid and 0.1 mg vitamin B12. The 1000x supplementation mix contained, per liter, 2.0 mg biotin, 5.0 mg thiamine-HCl, 2.0 mg folic acid, 0.1 mg vitamin B12, 127 μg/L spermidine, 174 μg/L spermine tetrahydrochloride and 0.7 mg hemin.

The aerobic cultures were inoculated to a starting OD_600_ of around 0.05, after an overnight culture on the base TMM medium. Growth was monitored through OD600 measurements, during growth at 60°, 200 rpm in baffled shake flasks. Anaerobic medium was prepared similarly to aerobic medium, but 1 μg/L resazurin was added to ensure complete anaerobic conditions. The medium was flushed with nitrogen gas prior to use. All anaerobic work was performed in an anaerobic chamber. Overnight cultures were run in anaerobic serum flasks at 60°C, 200 rpm and used to inoculate a microtiter plate to a final OD of 0.05 in 200 μL volume. After sealing, the OD_600_ was measured every 15 minutes for 10 hours in a Biotek Epoch2 microplate spectrophotometer, placed in an anaerobic chamber (run at 60°C, with linear shaking).

## 5. Abbreviations Metabolites

13dpg: 3-Phosphoglyceroyl phosphate
2pg: 2-phosphoglycerate
3pg: 3-phosphoglycerate
6pgl: 6-phosphogluconolactone
ac: acetate
acald: acetaldehyde
accoa: acetyl-CoA
actp: acetyl phosphate
akg: α-ketoglutarate
asp: aspartate
cit: citrate
dhp: dihydroxyacetone phosphate
e4p: erythrose 4-phosphate
etoh: ethanol (etoh)
f6p: fructose-6-phosphate
fdp: fructose 1,6-bisphosphate
for: formate
fum: fumarate
g3p: glyceraldehyde-3-phosphate
g6p: glucose-6-phosphate
glc: glucose
gly: glycine
icit: iso-citrate
lac: L-lactate
mal: malate
oaa: oxaloacetate
pep: phosphoenolpyruvate
phe: phenylalanine
pyr: pyruvate
r5p: ribose-5-phosphate
ru5p: ribulose-5-phosphate
s7p: sedoheptulose 7-phosphate
ser: serine
succ: succinate
succoa: succinyl-CoA
udpg: uridine diphosphate glucose
xu5p: xylulose-5-phosphate

## Reactions

ACKr: Acetate kinase
ACONTa: Aconitase
ALCD2x: Alcohol dehydrogenase
ASPTA: Aspartate transaminase
CS: Citrate synthase
DDPA: 3-deoxy-D-arabino-heptulosonate 7-phosphate synthetase
ENO: Enolase
FBA: Fructose-bisphosphate aldolase
FUM: fumarate hydratase
G6PDH2r: Glucose 6-phosphate dehydrogenase
GALUi: UTP-glucose-1-phosphate uridylyltransferase
GHMT: Glycine hydroxymethyltransferase
GLCtpts: Glucose phosphotransferase transporter
GLUSy: Glutamate synthase
ICL: Isocitrate lyase
LDH_L: L-lactate dehydrogenase
MDH: Malate dehydrogenase
PC: Pyruvate carboxylase
PDH: Pyruvate dehydrogenase
PFL: Pyruvate formate lyase
PGCD: Phosphoglycerate dehydrogenase
PGI: Glucose-6-phosphate isomerase
PGK: Phosphoglycerate kinase
PPCK: Phosphoenolpyruvate carboxykinase
PRPPS: Phosphoribosylpyrophosphate synthetase
PSCVT: 3-phosphoshikimate 1-carboxyvinyltransferase
SUCDi: Succinate dehydrogenase
SUCOAS: Succinyl-CoA synthetase
TALA: Transaldolase
TKT1: Transketolase
TKT2: Transketolase

## Availability of data and materials

The metabolic model, scripts and corresponding datasets generated during the study are all freely available, under an Apache 2.0 license, at the GitHub repository: https://github.com/biosustain/p-thermo/releases/v1.0.

## Acknowledgements

We thank Dr. Shyam Maskapalli for help with biomass composition analysis.

## Supplementary Figures

**Supplementary figure 1:**
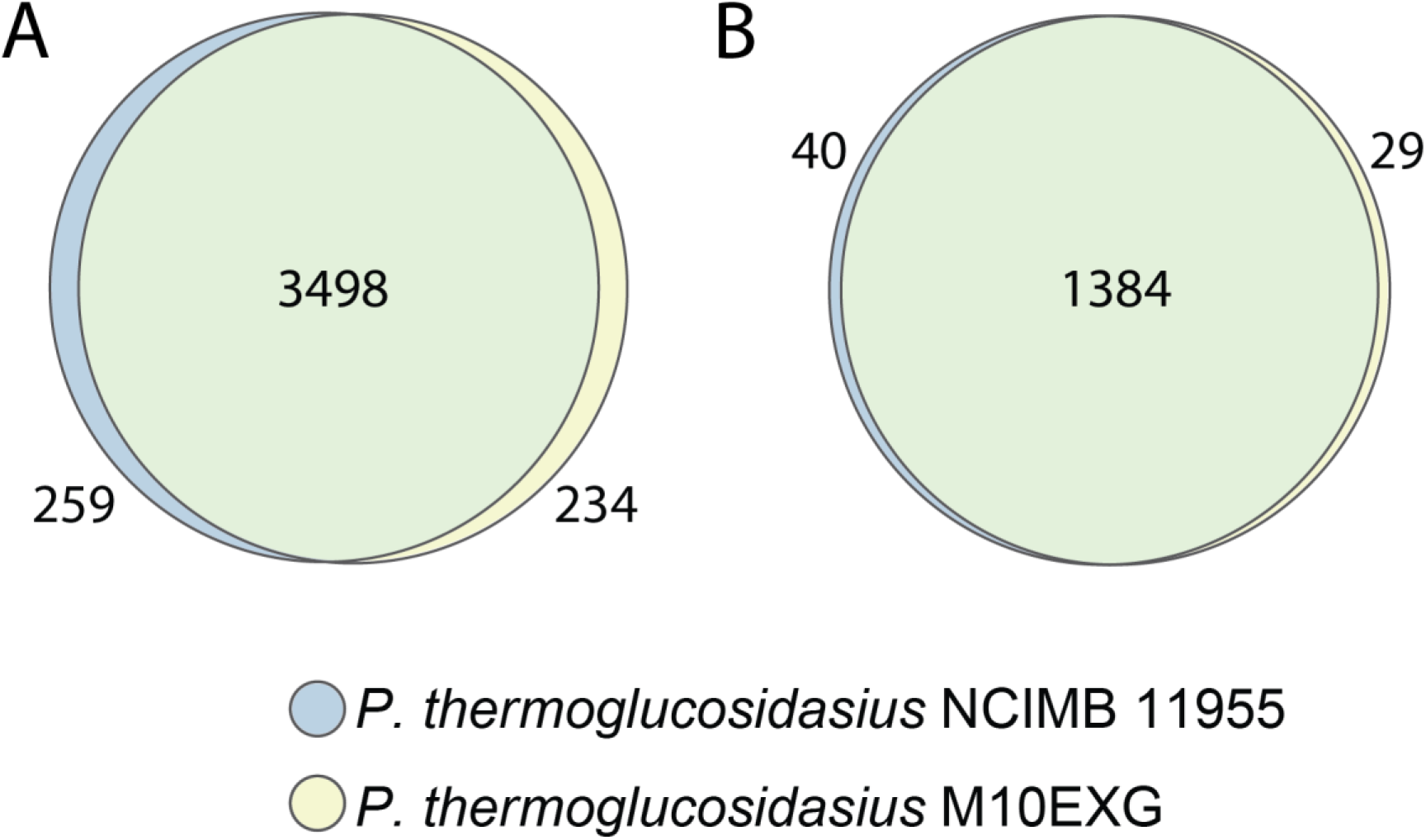
Whole proteome comparison between *P. thermoglucosidasius* NCIMB 11955 and *P. thermoglucosidaius M10EXG*, for all ORFs (A) and when filtered for metabolic genes (B, i.e. genes with an EC number associated to them). Supplementary tables 4 and 5 list the unique metabolic ORFs between the two strains.

**Supplementary figure 2:**
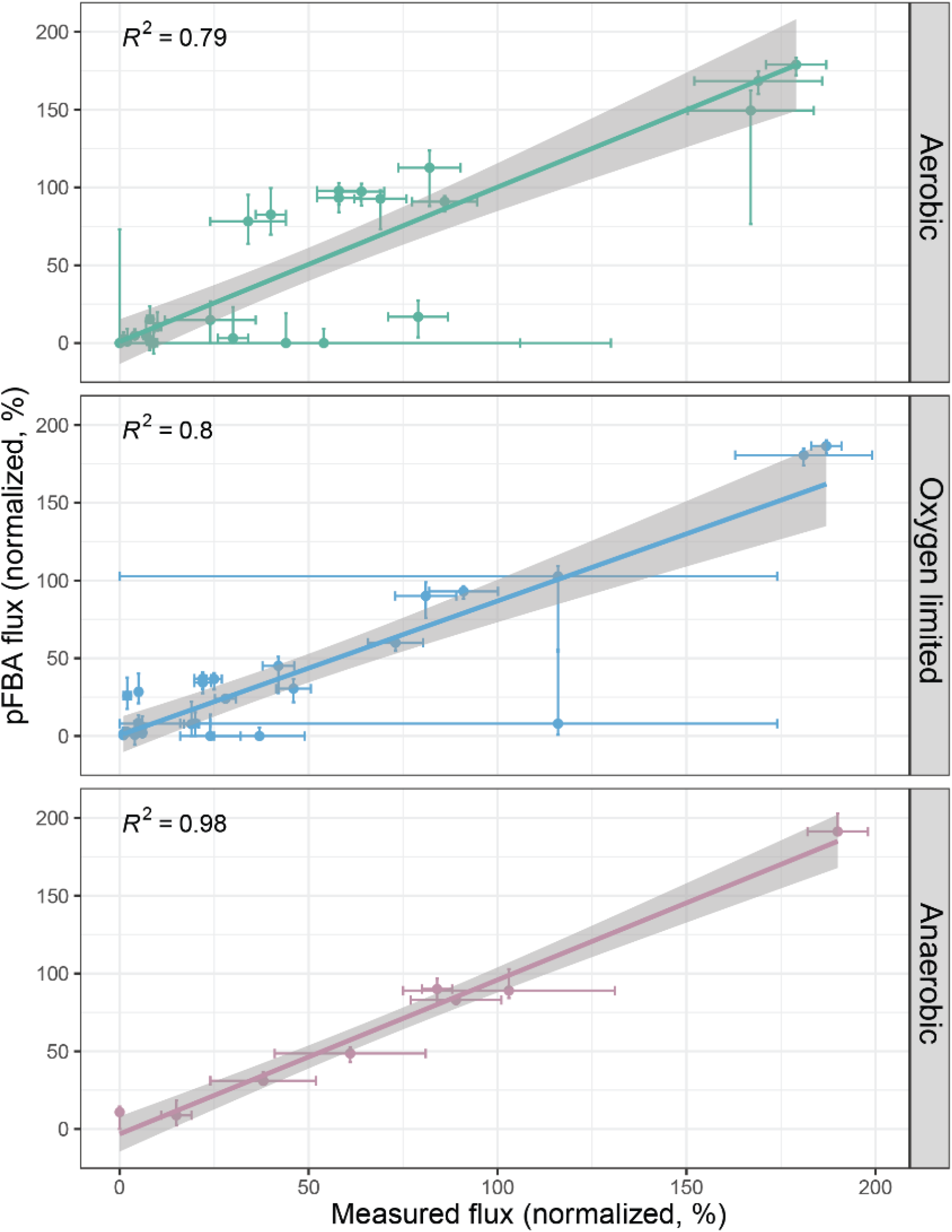
Correlation between pFBA analysis of the model and experimentally derived data^7^, normalized to glucose uptake rate, in aerobic, oxygen limited and anaerobic conditions. Errors for measured fluxes and variability in pFBA fluxes are shown. A linear fit has been applied to assay correlation, with the R value indicated per condition.

**Supplementary figure 3:**
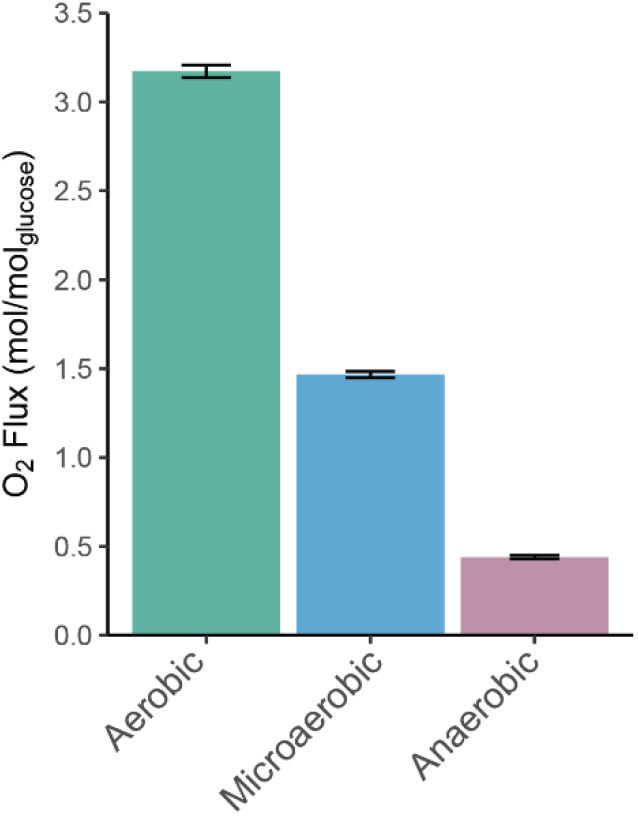
Predicted oxygen consumption rates for the three conditions, when measured exchange rates of fermentation products were fit to the model.

**Supplementary figure 4:**
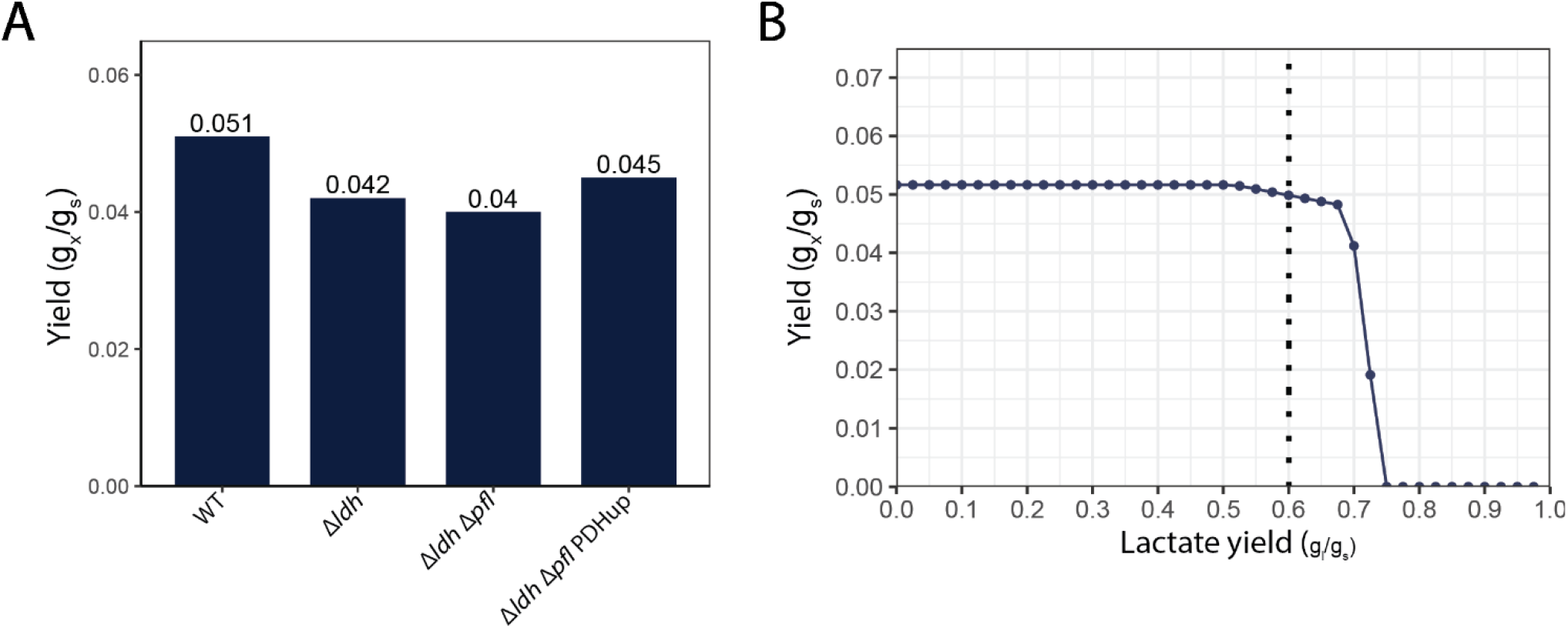
A) result of computing predicted *in silico* biomass yield, when measured exchange rates, carbon uptake rates and genetic manipulations (i.e. knockouts) have been introduced. B) Effect of lactate production on biomass yield when all other measured exchange rates are fixed for the WT strain. The dotted line indicates the measured lactate production in these conditions.

**Supplementary figure 5:**
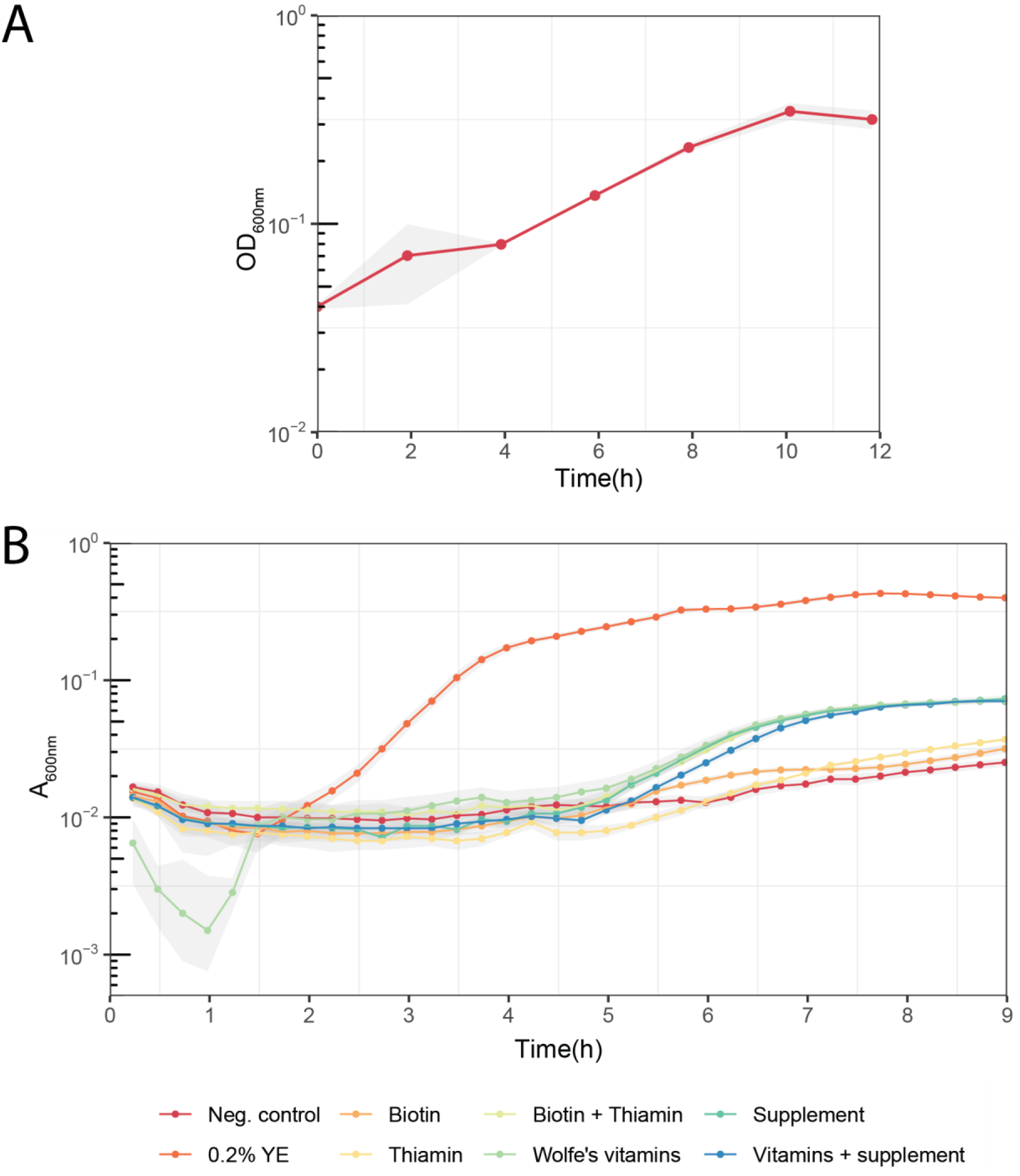
A) Aerobic shake flask experiment of *P. thermoglucosidasius* NCIMB 11955 on TMM base medium. Shaded erea shows standard devaition between three biological replicates. B) Anaerobic growth curves of *P. thermoglucosidasius* NCIMB 11955 grown on base TMM supplemented with various nutrients. Experiment was performed in a microtiter plate reader, and shaded area represents standard deviation of measurements over quadruplicates. (YE = yeast extract)

## Supplementary Tables

**Supplementary Table 1:** Stoichiometry of the biomass reaction in *p-thermo*.

|  | Compound | mmol/gDCW |
| --- | --- | --- |
| Proteins | Ala | 0.5142 |
|  | Arg | 0.1831 |
|  | Asp | 0.2219 |
|  | Asn | 0.2219 |
|  | Cys | 0.1079 |
|  | Glu | 0.322 |
|  | Gln | 0.322 |
|  | Gly | 0.4077 |
|  | His | 0.0778 |
|  | Ile | 0.2728 |
|  | Leu | 0.3475 |
|  | Lys | 0.3172 |
|  | Met | 0.0832 |
|  | Phe | 0.1451 |
|  | Pro | 0.1665 |
|  | Ser | 0.1811 |
|  | Thr | 0.2688 |
|  | Trp | 0.1026 |
|  | Tyr | 0.1036 |
|  | Val | 0.3761 |
| Nucleic Acids | *Ribonucleic acid (RNA)* |  |
|  | AMP | 0.1193 |
|  | CMP | 0.0915 |
|  | GMP | 0.0915 |
|  | UMP | 0.1193 |
|  | *Deoxyribonucleic acid (DNA)* |  |
|  | dAMP | 0.0077 |
|  | dCMP | 0.0059 |
|  | dGMP | 0.0059 |
|  | dTMP | 0.0077 |
| Lipids | *Phospholipids* |  |
|  | PE | 0.0290 |
|  | PG | 0.0277 |
|  | CLPN | 0.0296 |
| Carbohydrates |  |  |
|  | D-Fructose | 0.1048 |
|  | GDP-Mannose | 0.0057 |
|  | UDP-D-Galactose | 0.1895 |
|  | UDP-D-Xylose | 0.1825 |
|  | UDP-D-Glucose | 0.0096 |
|  | UDP-L-Arabinose | 0.0407 |
| Salts | Phosphorous | 0.0420 |
|  | Calcium | 0.0028 |
|  | Potassium | 0.6323 |
|  | Magnesium | 0.0875 |
|  | Iron | 0.0304 |
| Vitamins & cofactors | Thiamin B1 | 0.0002 |
|  | Riboflavin B2 | 0.0002 |
|  | Vitamin B12 | 0.0002 |
|  | Vitamin B6 | 0.0002 |
|  | 10-Formyltetrahydrofolate | 0.0004 |
|  | Biotin | 1.79E-06 |
|  | Bacillithiol | 6.23E-05 |
|  | Bacillithiol disulfide | 1.56E-07 |
|  | Chorismate | 0.0002 |
|  | FAD | 0.0002 |
|  | FMN | 0.0002 |
|  | Heme | 0.0002 |
|  | Proto-heme | 0.0002 |
|  | Siroheme | 0.0002 |
|  | Menaquinol | 0.0003 |
|  | NAD+ | 0.0145 |
|  | NADH | 0.0267 |
|  | NADP | 0.0042 |
|  | NADPH | 0.0027 |
|  | S-Adenosyl-L-methionine | 0.0002 |
|  | Spermidine | 0.0011 |
|  | Spermine | 0.0025 |
|  | Putrescine | 0.0054 |
|  | CoA | 0.0002 |
| Intracellular Metabolites | Acetyl-CoA | 0.0002 |
|  | Succinyl-CoA | 8.75E-05 |
| Energy Requirement | ADP | - 152.28 |
|  | Pi | -152.27 |
|  | Ppi | 0.0011 |
|  | H | - 152.28 |
|  | H2O | 152.28 |
|  | ATP | 152.31 |

**Supplementary table 2:** Estimation of polymerization energy needed to form biomass from the different monomer classes present in the biomass reaction. This energy fraction constitutes part of the growth associated maintenance that is present in the biomass reaction. Polymerization energy per molecule was obtained from literature^64^.

|  | Cellular content<br>(w/w) | Polymerization energy |  |
| --- | --- | --- | --- |
|  |  | Per molecule<br>(mmATP/g polymer) | Total (mmol<br>ATP/gcell) |
| Protein | 0.52 | 37.7 | 19.604 |
| Carbohydrates | 0.1 | 12.8 | 1.28 |
| RNA | 0.16 | 26 | 4.16 |
| DNA | 0.1 | 26 | 2.6 |
| Lipids | 0.09 | 25.6 | 2.304 |
| Sum |  |  | 29.948 |

**Supplementary table 3:** Overview of an analyses of filtering the unique ORFs detected in the genome analyses between two *(Para)geobacillus* strains, when various levels of filtering are applied to elucidate how many reactions would be unique in models made of each strain, and finally which would be connected to any pre-existing metabolites in the network. Supplementary table 4 and 5 highlight the unique metabolic ORFs identified.

|  | *P. thermoglucosidasius*<br>NCIMB 11955 | *P. thermoglucosiodasius*<br>M10EXG |
| --- | --- | --- |
| Unique ORFs | 259 | 234 |
| Unique metabolic ORFs<sup>1</sup> | 40 | 29 |
| Unique ORFs as reactions<sup>2</sup> | 18 | 13 |
| Unique reactions<sup>3</sup> | 11 | 12 |
| Connected unique reactions<sup>4</sup> | 8 | 6 |

<sup>1</sup> Unique metabolic ORFs are ORFs with an E.C. code associated to them
<sup>2</sup> Subset of unique metabolic ORFs that would actually be captured as a reaction in the model
<sup>3</sup> Subset of unique ORFs that would cause new reactions to the model
<sup>4</sup> Subset of unique reactions that are connected by a main metabolite to one or more pre-existing metabolites in the model. Note, all these reactions are still blocked (i.e. no two main metabolites found in model).

**Supplementary table 4:**
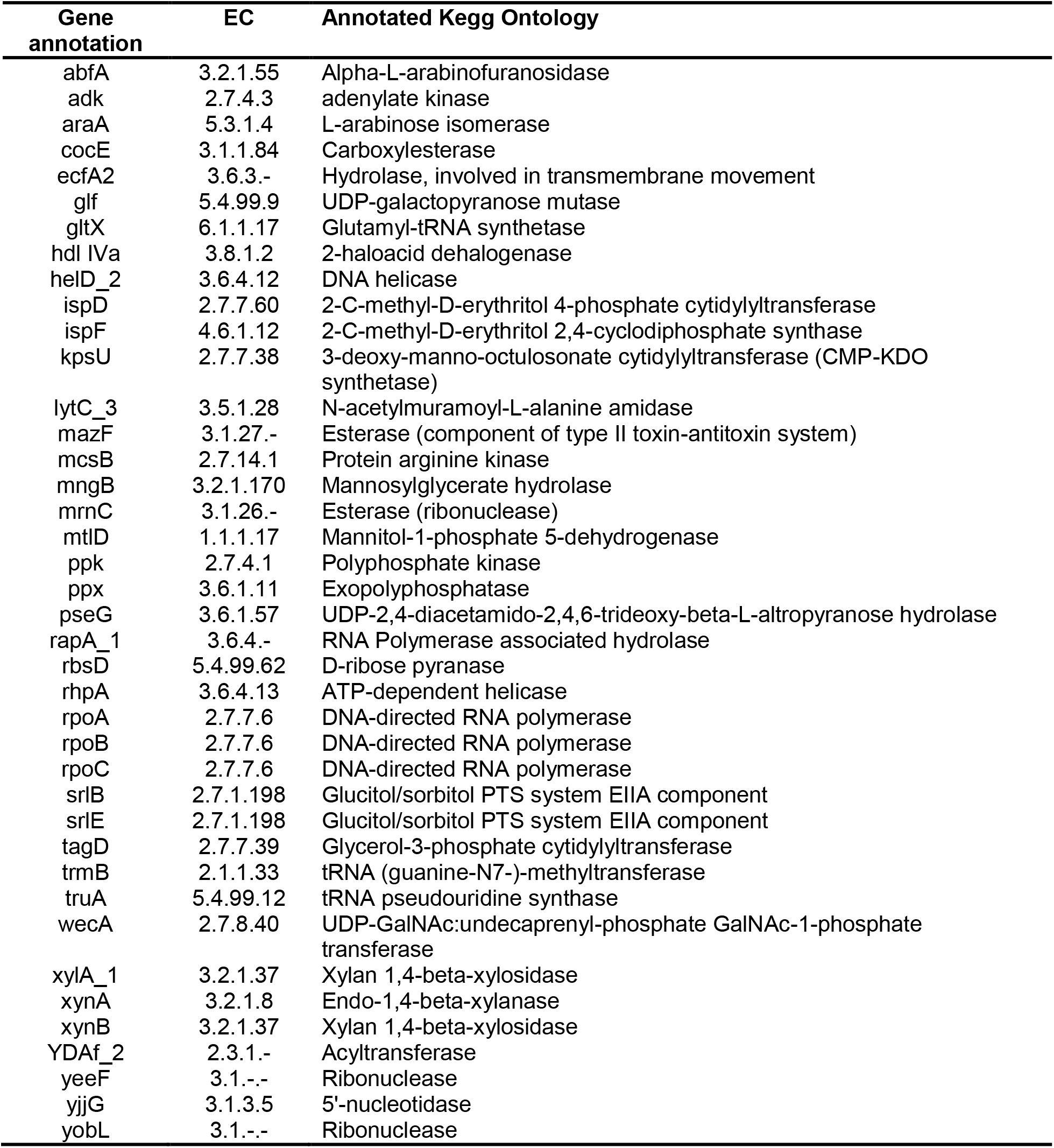
Metabolic ORFs unique to *P. thermoglucosidasius* NCIMB 11955, detected in the genome comparison.

**Supplementary table 5:** Metabolic ORFs unique to *P. thermoglucosidasius* M10EXG, detected in the genome comparison. Interestingly, it appears the M10EXG strain encodes a complete myo-inositol utilizing operon (*iol*), where some components are lacking in NCIMB 11955^103^. Data shows NCIMB 11955 indeed lacks the capability to grow on inositol^51^, but data on M10EXG is lacking.

| Gene annotation | EC | Annotated Kegg Ontology |
| --- | --- | --- |
| cocE | 3.1.1.84 | Carboxylesterase |
| cwlK | 3.4.-.- | Peptidase |
| cynS | 4.2.1.104 | Cyanate lyase |
| derK | 2.7.1.210 | D-erythrulose 4-kinase |
| dhaL | 2.7.-.- | Phosphotransferase |
| fdtA | 5.3.2.3 | TDP-4-oxo-6-deoxy-alpha-D-glucose-3,4-oxoisomerase |
| hsdM_1 | 2.1.1.72 | Site-specific DNA-methyltransferase (adenine-specific) |
| hsdM_2 | 2.1.1.72 | Site-specific DNA-methyltransferase (adenine-specific) |
| hsdR_1 | 3.1.21.3 | Type I restriction enzyme |
| hsdR_2 | 3.1.21.3 | Type I restriction enzyme |
| iolB | 5.3.1.30 | 5-deoxy-glucuronate isomerase |
| iolE | 4.2.1.44 | Inosose dehydratase |
| iolI | 5.3.99.11 | 2-keto-myo-inositol isomerase |
| iolX | 1.1.1.370 | Scyllo-inositol 2-dehydrogenase (NAD+) |
| lhgO | 1.1.3.15 | Glycolate oxidase |
| lipA | 2.8.1.8 | Lipoyl synthase |
| lsrF | 2.3.1.245 | 3-hydroxy-5-phosphonooxypentane-2,4-dione thiolase |
| mutT4 | 3.6.1.- | Esterase |
| NA | 2.4.1.161 | Oligosaccharide 4-alpha-D-glucosyltransferase |
| radD | 3.6.4.12 | DNA repair helicase |
| recD | 3.1.11.5 | Exodeoxyribonuclease V |
| rfbC | 5.1.3.13 | dTDP-4-dehydrorhamnose 3,5-epimerase |
| sqhC | 4.2.1.137 | Sporulenol synthase |
| sunS | 2.4.1.- | Glycosyltransferase |
| tatD | 3.1.21.- | Endodeoxyribonucleases |
| uvrD1 | 3.6.4.12 | ATP-dependent DNA helicase |
| wapA_1 | 3.1.-.- | tRNA(Glu)-specific nuclease |
| wapA_2 | 3.1.-.- | tRNA(Glu)-specific nuclease |
| wapA_3 | 3.1.-.- | tRNA(Glu)-specific nuclease |

